# Topographic mapping as a basic principle of functional organization for visual and prefrontal functional connectivity

**DOI:** 10.1101/549345

**Authors:** Jonathan F. O’Rawe, Hoi-Chung Leung

## Abstract

The organization of region-to-region functional connectivity has major implications for understanding information transfer and transformation between brain regions. We extended connective field mapping methodology to 3-dimensional anatomical space to derive estimates of cortico-cortical functional organization. Using multiple publicly available human (both male and female) resting-state fMRI data samples for model testing and replication analysis, we have three main findings. First, we found that the functional connectivity between early visual regions maintained a topographic relationship along the anterior-posterior dimension, which corroborates previous research. Higher order visual regions showed a pattern of connectivity that supports convergence and biased sampling, which has implications for their receptive field properties. Second, we demonstrated that topographic organization is a fundamental aspect of functional connectivity across the entire cortex, with higher topographic connectivity between regions within a functional network than across networks. The principle gradient of topographic connectivity across the cortex resembled whole brain gradients found in previous work. Last but not least, we showed that the organization of higher order regions such as the lateral prefrontal cortex demonstrate functional gradients of linearity and convergence. These organizational features of the lateral prefrontal cortex predict task based activation patterns, particularly visual specialization and higher order rules. In sum, these findings suggest that topographic input is a fundamental motif of functional connectivity between cortical regions for information processing and transfer, with maintenance of topography potentially important for preserving the integrity of information from one region to another.

**Significance Statement:** Quantifying spatial patterns of region-to-region functional connectivity provides an avenue for testing theories of corticocortical information transformation and organization. This work demonstrates that this quantification is feasible not only in early visual cortex, but even in highly multimodal regions where spatial topography is less clear. Overall, we show that topographic relationships as a common motif functional connectivity across the cortex between regions within the same functional network and that analyzing the lateral prefrontal cortex in terms of topographic connectivity reveals organizational features that voxelwise connectivity analysis misses.

## 1. Introduction

One primary goal of systems neuroscience is to understand how integration occurs across cortical areas in order to support complex behavior (Cavada & Goldman-Rakic, 1991; Haber, Fudge, & McFarland, 2000; Hoover & Strick, 1993). As different brain regions display spatially specific patterns of long range and short range connections to their target regions (Kravitz, Saleem, Baker, & Mishkin, 2011; Kravitz, Saleem, Baker, Ungerleider, & Mishkin, 2013; Van Essen & Gallant, 1994), it is believed that the particular properties of brain circuitry must provide a foundation for brain function and information processing, and that connectivity profiles from one region to another allows for the potential understanding of information transfer/transformation. Topographic, convergent, and divergent connectivity have been postulated to support different computational needs of a specific neural network (Thivierge & Marcus, 2007). For example, systematic convergent connectivity in the visual hierarchy has been shown to be responsible for object processing, from lines (Hubel & Wiesel, 1962) to complex objects (Tanaka, 1997), while divergent connectivity has been theorized in motor systems for complex feedback mechanisms, such as motor efference copies (Wolpert & Flanagan, 2001).

Neuroimaging studies of brain connectivity, while indirect, have the benefit of their exhaustive sampling across the entire anatomical space, which complement the interpretations from the sparse sampling of rigorous anatomical tracing in animals. This is evident in functional connectivity studies using resting state fMRI data, from which a low dimensional structure of large scale networks has been consistently observed as a key organizational feature of the cortex (Biswal et al., 2010; Di, Gohel, Kim, & Biswal, 2013; Fox et al., 2005; Yeo et al., 2011). Indeed, decomposing the whole brain into its fundamental gradients have revealed a structure for the organization of large scale networks, with the primary gradient trending from unimodal networks to multimodal networks (Margulies et al., 2016). However, despite the overwhelming focus on large scale brain organization, detailing the particular patterns of connectivity from one region to another remains a challenge.

Region-to-region topographic connectivity has been observed between early visual areas, which show retinotopic organization (Dumoulin & Wandell, 2008; Wang, Mruczek, Arcaro, & Kastner, 2015). By systematically seeding functional connectivity from known functional gradients, it has been shown that functional connectivity within a visual region, and between visual regions seem to follow along known eccentricity organization, along the anterior-posterior axis (Arcaro, Honey, Mruczek, Kastner, & Hasson, 2015). In other words, certain parts of a visual region that represent certain eccentricities are more likely to connect to the specific parts of other visual regions that represent similar eccentricities. While this type of systematic seeding is possible in areas with a known and measurable sensory topology, a more general model, such as connective field mapping (Haak et al., 2013), is useful for studying the connection topology of higher order regions. The previous connective field mapping technique fits 2 dimensional surface based population receptive fields (pRFs) using cortical signal from one region as the input to another region (Haak et al., 2013). This method has been used to model early visual areas, and demonstrate maintenance of retinotopic organization across these visual regions (Gravel et al., 2014; Haak et al., 2013).

It is unclear whether the linear pattern of connectivity observed in early visual cortex is dependent on a shared sensory topological organization, or whether it is present in any highly connected network. In particular, for multimodal regions such as the prefrontal cortex, while it is known that various subdivisions are closely connected, the exact functional organization remains controversial (Goldman-Rakic, 1999; Petrides, 2005; Ungerleider, Courtney, & Haxby, 1998). In theoretical models, spatial topography in connectivity is postulated to serve the function of ensuring the fidelity of information transfer across a network (Thivierge & Marcus, 2007). Topographic organization of connectivity also has the potential to generate complex abstract representations by virtue of topographic overlap (Tinsley, 2009). This suggests that abstract computations can be performed by biases in spatial connectivity and filtering between regions of a large scale network, in the same way that has been suggested in the visual system (Tinsley, 2009; Van Essen & Gallant, 1994).

In this study, we have modified and extended the connective field model, allowing for the estimation of topography of functional connectivity between regions using resting-state fMRI data in order to infer potential fidelity of information transfer between regions within and across unimodal and multimodal networks. This method fits a 3-D isotropic Gaussian to the connectivity pattern in the mapping region from individual voxels in the seed region (Fig. 1A). This fitting procedure produced four parameters, 3 location parameters describing the preferred locus of connectivity and a spread parameter, and the fitted Gaussian acts like an encoding model for timecourses of BOLD activity (Fig. 1B). We further estimated the degree of linear mapping from one region to another by performing Procrustes analysis. The degree to which these coordinates can be fit using a linear transformation provides the estimation of the linear mapping between them. We first examined the degree of topography connectivity in the visual system, as the literature is relatively established, and then examined whether it is simply a specific case of a general motif of brain functional connectivity across various networks, and finally used this approach to examine prefrontal cortex functional organization.

**Figure 1:**
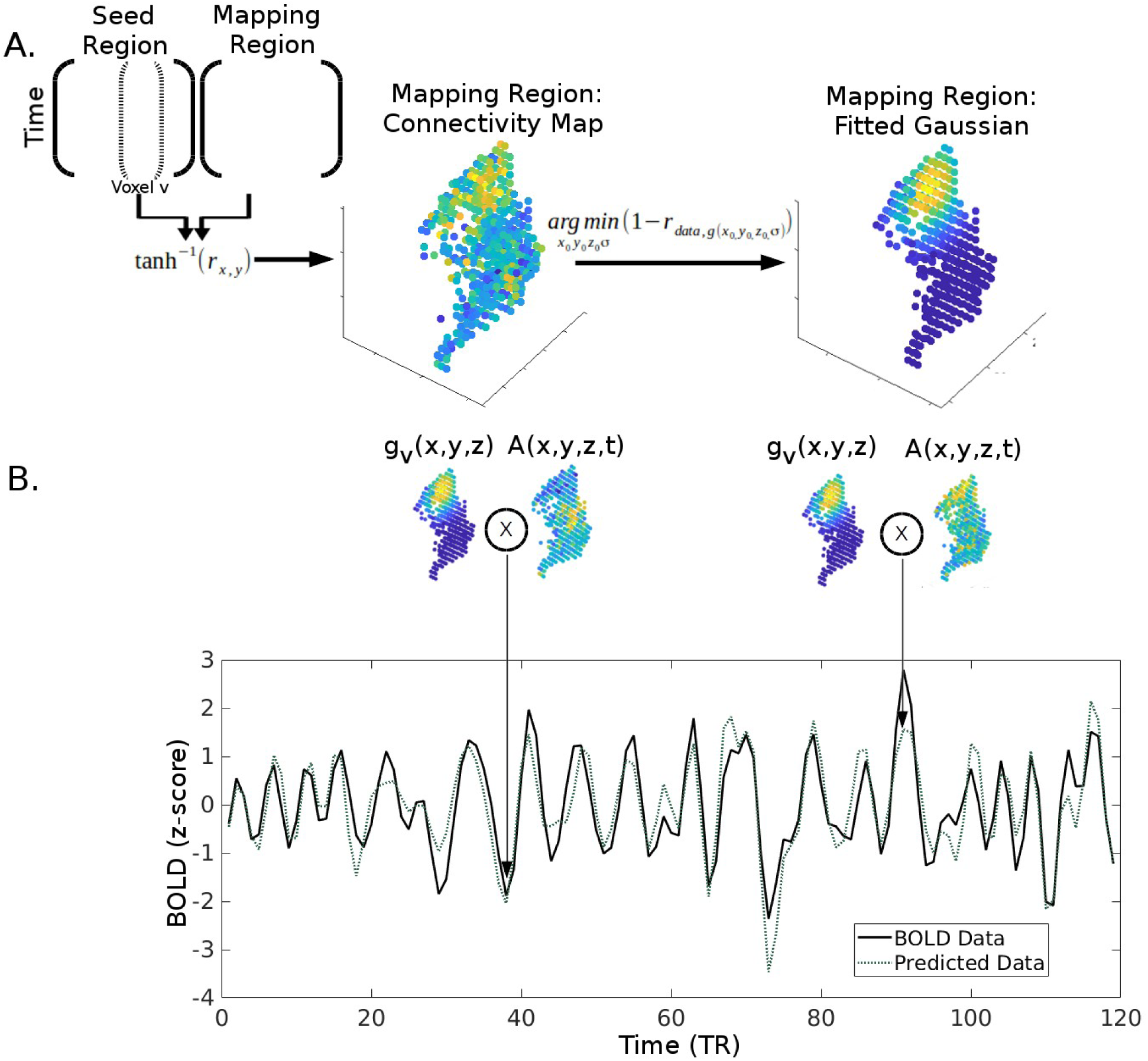
3D Connective Field Modeling. **(A)** Gaussian fitting procedure. Each seed voxel’s timecourse (dotted bracket inside the matrix on the left represents the timecourse of one voxel of the seed region) is correlated to every voxel’s timecourse in the mapping region and then Fisher’s z transformed. For each seed voxel, the corresponding 3-D Fisher’s Z matrix is then fitted to the 3 dimensional Gaussian model, by minimizing the correlation distance between the Fisher’s Z matrix and the Gaussian distribution. The fitted Gaussian parameters, location and standard deviation parameters, for each seed voxel are used in subsequent quantitative tests. **(B)** Time-series prediction model. For each time-point, the overlap between the actual activation in the mapping region [A(x,y,z,t)] and the fitted Gaussian distribution in the mapping region [g_v_(x,y,z)] for that particular seed voxel produce the expected level of activity in the corresponding seed voxel. The first arrow points to a period of low overlap, and thus a period of low expected level of activity, while the second arrow points to a period of high overlap, and thus a period of low expected level of activity. This procedure was used to evaluate the performance of the 3-D connective field model.

## 2. Materials and Methods

### 2.1 Subjects

We used three publicly available datasets: one sample of healthy controls with well validated resting state fMRI data, which is the Cambridge Buckner subset of the 1000 functional connectomes project (Biswal et al., 2010), and two smaller samples more well suited for test-retest reliability studies. The first small sample being the Intrinsic Brain Activity Test Retest (IBATRT) dataset (Zuo et al., 2014), and the second being the Midnight Scan Club (MSC) dataset (Gordon et al., 2017).

For the Cambridge Buckner data sample there were a total of 198 subjects (123 female), ages 18-30 (M = 21.03, SD = 2.31), with all subjects included in the final analysis. For the IBATRT data sample, there were a total of 36 subjects (18 female) ages 19-48 (M = 27.33, SD = 7.86) with two sessions and each with two runs. Four IBATRT subjects were excluded due to excessive motion in at least one of the four runs, leaving 32 subjects with data in the two runs of Session 1 (16 female; ages 19-48, M = 26.03, SD = 7.24). For the MSC dataset, there were a total of 10 subjects (5 female), ages 24-34 (M = 29.1, SD = 3.3), and all included in the final analysis.

### 2.2 fMRI Parameters

Cambridge Buckner data (Siemens 3T Trim Trio): T1-weighted images were collected with MPRAGE with the following image parameters: slices = 192, matrix size = 144 x 192, voxel resolution = 1.20 x 1.00 x 1.33 mm^3^. Resting state fMRI data were T2*-weighted images acquired using EPI with the following parameters: 47 interleaved axial slices, TR = 3000 ms, voxel resolution = 3.0 x 3.0 x 3.0 mm^3^ (119 volumes).

IBATRT data (Siemens 3T Trim Trio): T1 weighted images were collected with MPRAGE with the following image parameters: slices = 176, matrix size = 256 x 256, voxel resolution = 1.0 x 1.0 x 1.0 mm^3^. Resting state fMRI data were T2*-weighted images acquired using EPI with the following parameters: 29 ascending axial slices, slice gap = 0.36 mm, TR = 1750 ms, voxel resolution = 3.4 x 3.4 x 3.6 mm^3^ (343 volumes each run). While there were up to 4 runs across 2 sessions for subjects, we only utilized the two runs in the first session.

MSC Data (Siemens 3T Trim Trio): Four T1 weighted images were collected: slices = 224, voxel resolution = 0.8 x 0.8 x 0.8 mm^3^. Four T2 weighted images: 224 slices, voxel resolution 0.8 x0.8 x 0.8 mm^3^. Resting state fMRI were T2* weighted images acquired using EPI: 36 interleaved axial slices, TR = 2200 ms, voxel resolution = 4 x 4 x 4 mm^3^ (818 volumes each session). A gradient echo field map was collected with the same parameters as the BOLD EPI images for each session. There were 10 sessions for each subject, all utilized in the analysis.

### 2.3 Image Preprocessing

For each individual in the Cambridge Buckner and IBATRT data samples, preprocessing was performed utilizing SPM12 (http://www.fil.ion.ucl.ac.uk/spm/software/spm12/). The functional images were first corrected for slice timing, and then realigned to the middle volume according to a 6 parameter rigid body transformation. Structural images were coregistered with the mean functional image, segmented, and then normalized to the MNI template using both linear and nonlinear transformations. Functional images were normalized utilizing the same parameters as the structural normalization.

Further preprocessing was performed following the standard procedures of resting-state fMRI analysis either using CONN (Whitfield-Gabrieli & Nieto-Castanon, 2012) or custom Matlab (2015b) scripts. A nuisance regression was constructed with the following confounding variables: 6 motion parameters up to their second derivatives, scans with evidence of excessive motion (Framewise Displacement [FD] > .5 or DVARS > 5), session onset, estimated physiological signal generated through aCompCor (a temporal PCA of the white matter and CSF voxels with the number of components included determined individually on the basis of a Monte Carlo null model (Behzadi, Restom, Liau, & Liu, 2007)), and a linear drift component. For the Cambridge Buckner data the residuals from the nuisance regression were filtered utilizing a bandpass between the frequencies of 0.008 and 0.09 Hz, while for the IBATRT data, the band-pass filtering and nuisance regression were done simultaneously (Hallquist, Hwang, & Luna, 2013). Finally, the resultant data were despiked using a tangent squashing function.

For the MSC data, we utilized their release of preprocessed data which includes slice intensity correction, mode 1000 intensity normalization, realignment, transformation into Talairach space and FSL’s field distortion correction (Smith et al., 2004). These data also came with additional preprocessing for resting-state analysis. Censored volumes were determined by an FD threshold of 0.2 mm. The data were first demeaned and detrended, then a nuisance regression was performed removing the following factors, with censored volumes ignored: global signal, white matter mean signal, CSF mean signal, and motion regressors with the full Volterra expansion. The data were then interpolated across the censored volumes using least squares spectral estimation followed by band-pass filtering between 0.009 and 0.08 Hz. The censored volumes were removed in the final resultant data samples for analysis (see Gordon et al., 2017 for full description of processing pipeline).

### 2.4 Regions of Interest

The initial analyses of 3-D connective field mapping were conducted using the visual areas as regions of interest (ROIs). For the early visual areas, we utilized the probabilistic visual atlas (Wang et al., 2015), selecting all voxels with any probability of either dorsal or ventral visual cortex, with no overlap between the two ROIs (see Fig. 2A). To determine whether the finding from the overall visual cortex mask was a general property of early visual areas, we also defined more selective masks of the dorsal and ventral portions of V1, V2, and V3, each with probability greater than 30% to reduce overlap between the masks. We then applied the same connective field mapping analyses across each visual region’s dorsal and ventral portions.

**Figure 2:**
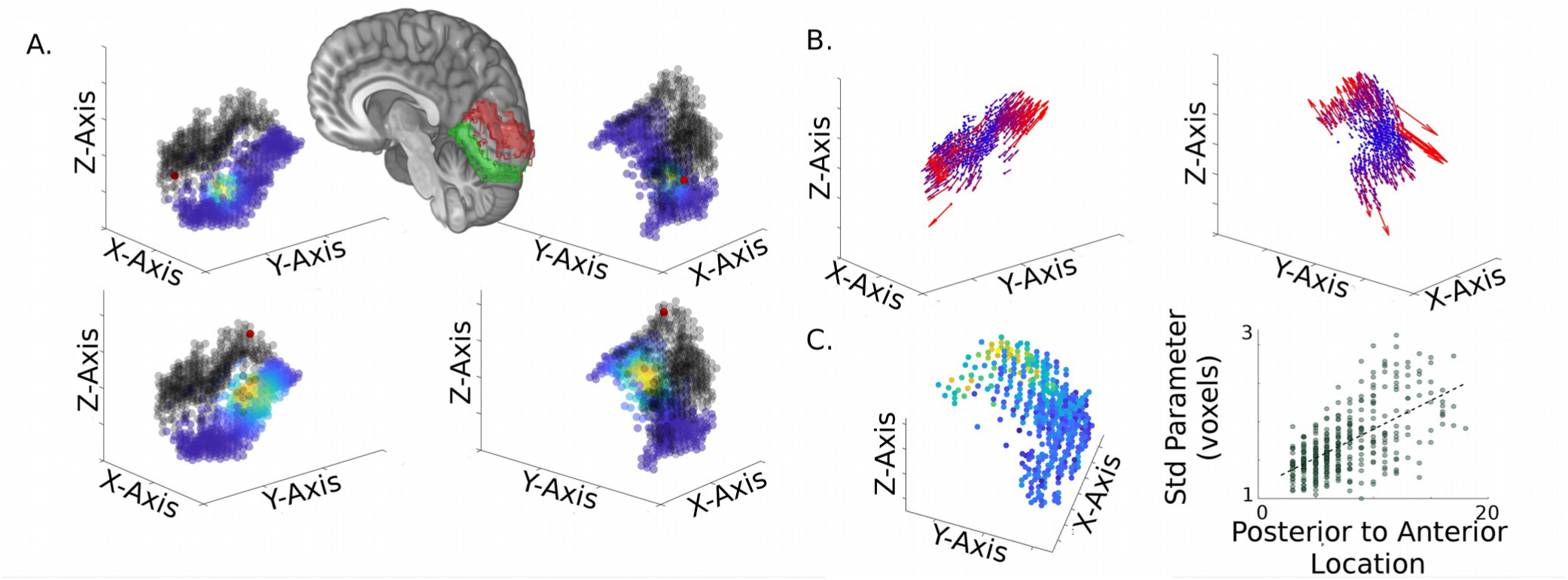
Group results of the 3-D connective field modeling in dorsal and ventral visual cortex using the Cambridge Buckner resting-state fMRI dataset. **(A)** Examples of the resulting Gaussian model are shown for two seed voxels (red dots), plotted in red among other voxels (black circles) in dorsal visual cortex (red masked area in the right hemisphere), and their associated group level Gaussian fit (color scale ranged from blue to yellow) in ventral visual cortex (green masked area). For full visualization, each two column shows two plots of the seed voxel’s fitted map rotated by 180 degrees from each other. Full animation can be found in Supplementary materials. **(B)** Preferred location of connectivity of each seed voxel is plotted as a vector with the origin in the seed location, and the vector direction showing the fitted x_0_ y_0_ and z_0_ parameters. The vectors are colored coded with blue to red representing low to high euclidean norm of the vector. **(C)** Group average of the Gaussian model’s standard deviation parameter. Left, each voxel within the seed region, dorsal visual cortex, is plotted with the data point’s color scaled by the standard deviation parameter of their connective field model. Cooler to warmer colors represent smaller to larger standard deviations. Right, this plot shows the correlation between the anterior-posterior extent of the seed voxel and their associated standard deviation parameter, suggesting that standard deviation increases along more anterior voxels.

In an additional analysis, we created slightly more lenient V1 mask selecting any voxels that have a greater than 15% probability of being V1. We used this V1 ROI as a mapping region for the right fusiform face area (FFA) and the right parahippocampal place area (PPA), both FFA and PPA were derived using Neurosynth meta-analyses, with “faces” and “place” as keywords, respectively (Yarkoni, Poldrack, Nichols, Van Essen, & Wager, 2011).

For the whole brain topographic connectivity analysis, we made use of a previously published whole brain parcellation atlas of 400 regions. The parcels were generated by maximizing correlations within a parcel while minimizing local changes in correlations within a parcel (Schaefer et al., 2018). For time and computational feasibility, we only used the 200 parcels of the right hemisphere.

### 2.5 3-D Connective field mapping and model parameters

The 3-D connective field mapping analysis has been released in the form of a MATLAB toolbox, with a simple GUI wrapper to help guide analysis (www.nitrc.org/projects/r2r_prf/).

For both the “seed” and “mapping” regions, the mean timecourse of the ROI was subtracted from each voxel’s timecourse, so as to reduce the influence of correlation at the mean signal level, though in practice the results are similar with or without removing the means. Then, for each voxel within the “seed” region, we calculated its correlation with every voxel within the “mapping” region and transformed the resulting correlation values using Fisher’s Z transformation. Each distribution of Fisher’s Z values (for each seed voxel) was fitted with a 3-D isotropic Gaussian distribution (Fig. 1A):

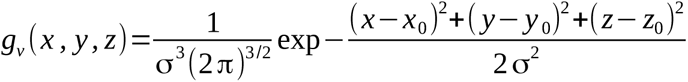

Initial values for the x_0_, y_0_, and z_0_ parameters during the optimization procedure were the x, y, and z locations of the maximum Fisher’s Z value, and an arbitrarily small number (floating point epsilon) for the standard deviation. The optimization procedure had a lower bounds of the smallest location parameters within the mapping region and the floating point epsilon for the standard deviation, while the upper bounds were the maximum location parameters and the smallest range of the 3 dimensions for the standard deviation. The parameters of the Gaussian were estimated by minimizing the correlation distance between the Fisher’s Z values and the Gaussian probability density function using the interior-point algorithm (Byrd, Gilbert, & Nocedal, 2000), implemented by MATLAB’s fmincon function (MathWorks Inc, 1995-2019). Predicted time-series for each voxel v in the seed region were estimated as the degree of overlap between the estimated Gaussian function and the activation pattern of the mapping region for each time-point t (Fig. 1B):

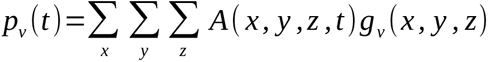

Both the predicted time-series and the actual data time-series were z-scored to remove any arbitrary scaling factors. Fit was assessed by comparing the distribution of correlations between the each voxel’s fitted model and the true data to the distribution of correlation between each voxel’s true data and every other voxel’s fitted model.

Topographic organization of functional connectivity between the seed and mapping region was estimated as rank linearity: estimated using the Procrustes method by fitting 3 space parameters of the Gaussian model to the 3 rank-ordered space parameters of the seed locations (Kendall, 1989). The three dimensional estimated parameters from the 3-D connective field model where first rank ordered, and then a transformation matrix with translation, rotation, and scale components was estimated that minimizes the normalized sums of squares error (SSE) between the rank ordered parameters and their spatial origin. The complement of the normalized SSE between the transformed parameters and the actual seed locations was used as a measure of the degree of topographic connectivity between two regions. We evaluated this linearity on a binomial distribution with respect to the independent observations within the seed region, given the observed smoothness of each subject’s BOLD data (see Supplemental Materials and Fig. S1-2 for full reasoning and validation through simulations). We performed statistical analysis on the MSC data, as reliable estimates were possible at the individual subject and individual session level. This was done to demonstrate the validity of the group level observations in the Cambridge-Buckner data sample. Further, vector fields were constructed to visualize the connectivity organization between two regions, with each vector originating at each seed voxel, and the orientation and length of the vector determined by the centered Gaussian parameters, its preferred location in the mapping region in relation to the center of mass of the mapping region (e.g., see Fig. 2B).

However, cortico-cortical relationships are likely to be obfuscated in 3D space due to cortical folding. While previous attempts to examine this have aligned data to the cortical surface to avoid this problem, we did not use cortical sheets for several reasons. First and foremost, since the data is acquired in volumetric space, most operations aligning to a cortical surface model results in additional smoothing to the data (e.g. through averaging). In addition, unfolding the cortical sheet is an arbitrary process, with distortions added in different forms depending how it is done. Therefore, constructing objective 2D coordinates is intractable. Our measures of topography provided novel information in the original space, but are likely influenced by the different folding patterns of the cortex to some extent. Therefore, we also visualize the relationship between regions by constructing a vector field. The vector field allows the researcher to observe when spatial relationships from one cortical region conform to the cortical surface, as the vector field should bend along sulci and gyri (see Fig. S3 for an example in a single subject, and group vector field on how vectors are distorted when constrained to the cortical sheet).

In addition to topographic connectivity, we also estimated convergence of functional connectivity. Convergence is the logical counterpart to topographic connectivity, and was calculated using a 3-D form of the Kolmogorov-Smirnov (KS) test (Fasano & Franceschini, 1987). The convergence metric was defined as the deviance of the Gaussian parameters from uniform distribution across the voxels in the mapping region. This provides a relatively distribution insensitive measure of convergence (Fasano & Franceschini, 1987). To correct for regional differences in spatial correlations and number of voxels, we used the KS value corrected for the KS value at the 5% alpha level. The relationship between convergence factor and rank linearity is shown in Figure S4.

We assessed test-retest reliability using the IBATRT and MSC datasets. For the IBATRT dataset, we examined the spatial correlation between the patterns of parameters in run 1 and run 2, within individual subjects and across session means. For the MSC dataset, we examined potential effects of added scan time to the stability of the 3-D Gaussian parameters for individual subjects. We split the each subjects’ sessions into two groups, a test and a retest group. We varied the number of sessions in each group, with a maximum of 5 session per group, and calculated the test-retest reliability for each subject. We also performed 3-D connective field mapping for each subject with all their sessions to demonstrate the stability of the group effect at the level of individual subjects.

### 2.6 Examining connectivity profiles in higher order visual regions

To study how the patterns of connectivity from early visual cortex to higher order visual regions, such as fusiform face area (FFA) and parahippocampal place area (PPA) (see section 2.4 for ROI selection), may predict their complex response properties, we fit 3-D connective field models seeding from these areas, mapping into primary visual cortex. We compared convergence and topographic connectivity between PPA/ FFA and V1, and compared the model parameters to those of the intrinsic convergence and topographic connectivity within V1.

We expected both regions to have higher convergence than the relationships between dorsal and ventral V1. Given the observed retinotopy in PPA (Arcaro, McMains, Singer, & Kastner, 2009), and the relatively weaker observed retinotopy in FFA (Saygin & Sereno, 2008), we also expected PPA to demonstrate a higher amount of topographic connectivity and a lower amount of convergence with V1 relative to FFA connectivity relationships with V1.

### 2.7 Whole brain characterization of topographic functional connectivity

We performed 3-D connective field mapping on each unique pair between all possible pairs out of the 200 parcels in the right hemisphere of the Shaefer atlas for each subject in the MSC data. For each fitted pair, we calculated the rank linearity of connectivity using Procrustes analysis. We selected several seed regions to test whether topography is maintained through specific networks or generally across the brain. We selected several a priori networks and examined whether their topographic characteristics follow network like behavior, namely that their within network topographic connectivity is stronger than between network topographic connectivity.

As a higher order summary, and to compare with previous whole brain gradient decompositions (Margulies et al., 2016; Murray, Demirtaş, & Anticevic, 2018), we performed nonmetric multidimensional scaling on the rank linearity matrices from all parcel pairs to estimate the gradients of topographic connectivity across the right hemisphere. The far ends of these gradients were considered to represent which regions maintain linear information across the brain. We produced a 2 dimensional solution, on the basis of an elbow in the plot of the rank correlation of the produced distance and the observed distance matrices (Fig. S10A), and visualized the gradients across the brain.

### 2.8 Lateral Prefrontal Cortex Analysis

We selected the subset of parcels (see section 2.4) within the lateral prefrontal cortex, and performed a similar gradient decomposition analysis specifically on prefrontal functional connectivity topography. We estimated two gradients (see Fig. S10B for rational) of the lateral prefrontal cortex and then examined whether the discovered organization reflects principles of prefrontal cortex organization described in the literature (Goldman-Rakic, 1987; Koechlin, Ody, & Kouneiher, 2003). For comparison, we further examined the gradients resulting voxelwise functional connectivity (Fig. S11).

Each participant of the MSC data sample also went through three tasks in the scanner, an incidental memory task in which participants judged various characteristics of faces, scenes, and words, a motor task in which participants were instructed to move various parts of their body, and a mixed block task design in which participants had to either discriminate between nouns and verbs, or coherently and incoherently moving dots, with a cue on the onset and offset of each block (Gordon et al., 2017). We used two contrasts to examine sensory specialization (Memory: Face – Scene, Mixed: Visual – Semantic Discrimination) and one to examine abstract rules (Mixed: Discrimination [Visual and Semantic] - Cue effects [block onset/offset]).

In order to examine how the organizational features of the lateral prefrontal cortex observed in the resting state scans relate to spatial patterns of task activation, we constructed three multiple regressions, one for each contrast, predicting task activation from each topographic gradient and average convergence.

## 3. Results

### 3.1 Mapping functional connectivity in the visual cortex

To replicate previous findings from 2-D modeling on the surface (Gravel et al., 2014; Haak et al., 2013), we first used 3-D connective field modeling to show that dorsal and ventral portions of the visual cortex have a topographic organization in their functional connectivity. Figure 2A illustrates the group average 3-D connective field maps in the ventral visual cortex for two selected seed voxels in the dorsal visual cortex using the Cambridge Buckner dataset. An animation of all seed voxels and their corresponding modeled Gaussian maps is available in the supplemental materials. Fig. 2B summarizes these results as a vector field in the seed region, with a vector originated from each seed voxel displaying the location of its preferred connectivity as the relative displacement from the center of the mapping region (Fig. 2B). This vector field of the dorsal visual cortex revealed a strong connectivity mapping to the ventral visual cortex along the predicted anterior-posterior organization (Arcaro et al., 2015), which was also supported quantitatively by the high correlation between the seed’s y-location and the mean modeled y-location of the Gaussian fit in the mapping region, with r(394) = 0.71, p = 1.63 x 10^-62^ (r = 0.83, with outliers removed). The spread or standard deviation parameter of the Gaussian model also increased along the seed region’s y-axis (Fig. 2C left), which was supported quantitatively by the correlation between the seed’s y-location and the modeled standard deviation parameter, with r(394) = 0.61, p = 5.45 x 10^-42^ (Fig 2C right). This relationship was similarly evident between the dorsal and ventral portions of V1, V1, and V3, using a set of more restricted masks (Fig. S5). These results together suggested that the inter-regional connectivity in early visual areas mirror the organization of eccentricity representation, with region to region connectivity ordered spatially along similar eccentricity locations, and the spread of that connectivity potentially following the increase in receptive field sizes toward more anterior visual cortex. While the group averaged data can provide qualitative aspects of the organization of regions, the averaging procedure can introduce various biases (and average out individual differences to produce an optimistically simple output).

To quantify the organization at an individual subject level, we first quantified whether or not these measures are stable at the individual subject level data, and further we moved to a data sample that has much longer BOLD collections to ensure high quality estimations (Midnight Scan Club). The median test-retest reliability within subjects increases with more scan time per subject, with the 150 minutes in each split leading to a test-retest reliability of 0.81 (Fig. S7C). A vector field of dorsal-ventral connectivity mapping for the visual cortex was generated for each individual and each session, and the anterior-posterior relationship was observed in every subject (Fig. S8). Quantitatively, the correlation between y-location modeled Gaussian parameter and seed y-location was high: ranging form 0.5 to 0.72 across the 10 individuals. In addition, topographic connectivity estimates for the entire dorsal and ventral visual cortex for each MSC individual ranged between 13.77%-18.27% (all p_FWE_’s < .05). Similarly, linearity estimates of V1d→V1v were highly replicated in each MSC subject, ranging from 24.62% to 34.71% (only one session of MSC02 and one session of MSC08 > α_FWE_).

We also evaluated the model validity and stability in several ways. First, we examined the distribution of correlations between the predicted timecourses from the model and the actual BOLD timecourses (see Fig. 1B) for each voxel in the large dorsal visual cortex mask (Fig. S6, left). The distribution of r-values across all subjects and all their correspond timecourses when using the matching prediction was strictly positive, with a mean of 0.55, whereas the distribution of correlations produced by nonmatched pairs is much wider with a mean of close to zero (Fig. S6, right). Second, also evaluated the stability of the 3-D connective field model within individual subjects and across mean results for the dorsal and ventral visual cortex, using the IBATRT dataset. For each subject, the estimated y-location parameters from the first and second runs were correlated across voxels (Fig. S7A). The resulting distribution of r-values had a median of 0.60, demonstrating moderate to good reliability at the individual subject level. At the group level, the mean parameters for each run correlated across voxels strongly, with r = 0.98 (Fig. S7B).

### 3.2 Connectivity bias in higher-order visual areas

Higher order visual regions have been characterized as complex filters of early visual region information (Van Essen & Gallant, 1994), with this type of model shown to predict FFA activation during a task (Kay & Yeatman, 2017). We thus examined to what extent the pattern of region to region functional connectivity reflect this filtering process. Using the MSC data, we applied the 3-D connective field mapping method to model the pattern of right FFA connectivity with V1 and compared its connectivity profile with that of PPA, a region which is selective to stimuli that have more peripherally distinctive features. Due to initial poor r^2^ of the Gaussian fits, we combined the sessions into two distinct sessions, allowing for better estimation across each. In comparison to the PPA, the FFA showed more connectivity with posterior and lateral V1 (t[9] = −7.04, p = 6.04 x 10^-5^ and t[9] = 3.36, p = .008, respectively) (Fig 3A). This is consistent the FFA receiving information from V1 areas with more foveal receptive fields. For visualization, we plotted group averaged vectors from each seed region to V1. From these vector fields, FFA seems to share more convergent and less topographic connectvity with V1 (Fig. 3B). We quantified this in individual subjects and compared across regions, with V1d to V1v connectivity for reference. When compared to PPA, the FFA has both significantly higher convergence and significantly lower topographic connectivity with V1 (t(9) = 5.25, p = 5.27 x 10^-4^ and t(9) = −4.33, p < .002, respectively) (Fig. 3C). PPA, when compared to measures within V1, in turn, shows lower linearity and higher convergence.

**Figure 3:**
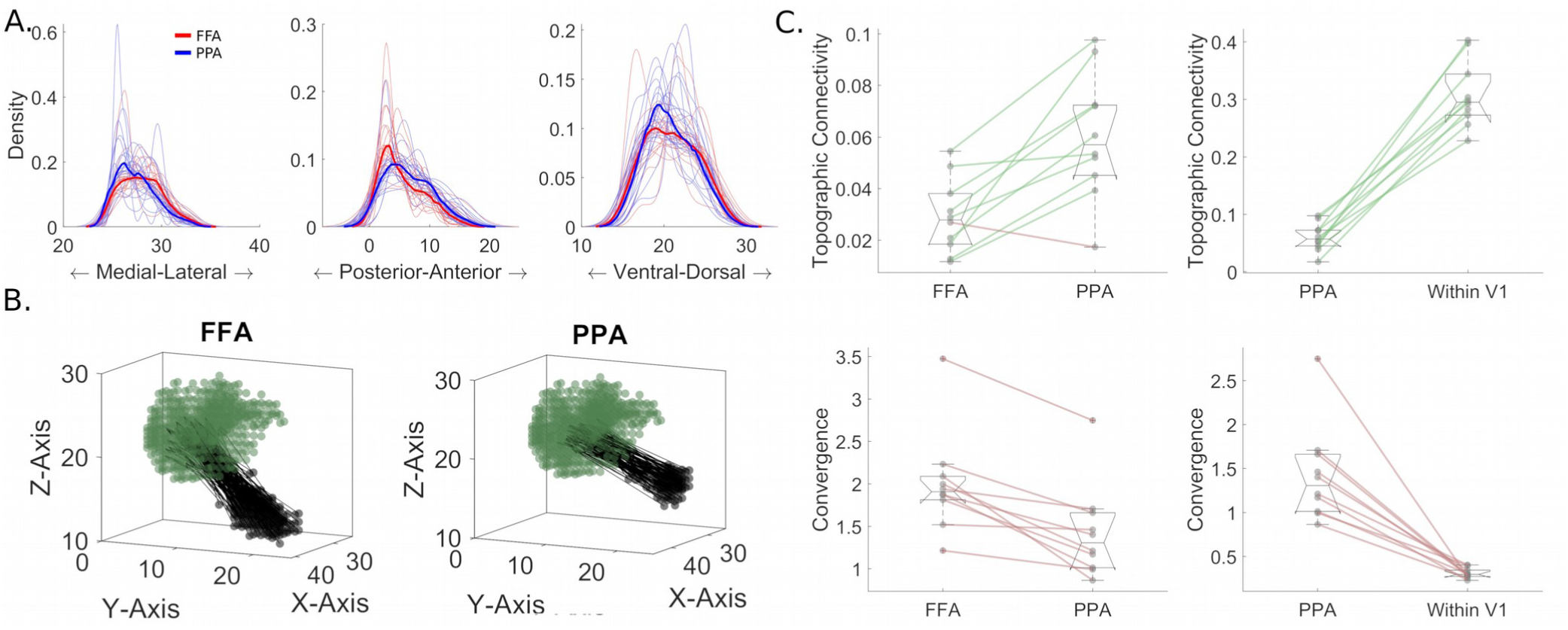
Pattern of functional connectivity between higher order visual regions and V1 corresponds well with their visual response properties from previous studies. **(A)** Compared to the PPA, FFA showed a more posteriorly located and narrower range of connectivity with V1. **(B)** Another representation of the biases of FFA to V1 connectivity (left) in comparison to PPA to V1 connectivity (right). The black points represent voxels within the seed region and the green points represent voxels within the mapping region, with the vectors connecting them represent the seed voxels’ preferred connectivity location along the mapping region. The mean vector fields provide information about quanitative features of the connectivity (such as the relatively more focal and mixing connectivity seen in FFA vs, PPA) but to quantify these results requires computation on individual subject vector fields. **(C)** Estimations of topographic connectivity show higher topography with V1 maintained in PPA, and higher convergence with V1 in FFA. Both of these metrics are shown next to dorsal and ventral V1 (within V1) estimates for reference.

### 3.3 Topographic connectivity organization across the cortex

While the early visual system demonstrated large topographic relationships in functional connectivity between regions, a fundamental question left unanswered is whether this is a special property of the early visual cortex or it is a general motif of cortico-cortical functional connectivity. To address this issue, we calculated the linearity of connectivity between all unique pairs of the 200 right hemisphere parcels in the Schaefer parcellation. We used the MSC dataset for this analysis to ensure stable parameter estimates, given the long scan time for each session. For each subject of the MSC dataset, the average percentage of connections that surpassed α = .05 threshold across all sessions and all pairs of parcels was 20.60% (range across subjects: 13.30%-25.69%). Note that the subject with the lowest number of significant topographic relationships, MSC08 (with 13.30% of connections surpassing .05 alpha), was identified by the original investigators to have a significant proportion of time in a drowsy state, affecting this individual’s BOLD data (Gordon et al., 2017). Therefore, on average, each brain region maintains a detectable topographic relationship with ∼20% of the rest of the brain, at a liberal threshold for detection. Eliminating parcels which likely do not follow the assumptions of our model (due to low number of resels) increases this average percentage (to 23.54%, ranging from 14.25%-29.64%).

Figure 4 illustrates the linear organization for three selected seed regions’ connectivity with other regions across the entire right hemisphere. The flatmaps show a clear segregation of linearity values within and across the different networks, exemplified by seeding from anterior and posterior visual cortex demonstrating stronger linearity with other visual areas, middle frontal gyrus demonstrating stronger linearity with lateral prefrontal and parietal regions, and lower linearity between regions from different networks. In Fig. S9, we present MSC01’s session by session seed rank linearity maps, presented with -log(p) values to demonstrate the stability of the statistical inference.

**Figure 4:**
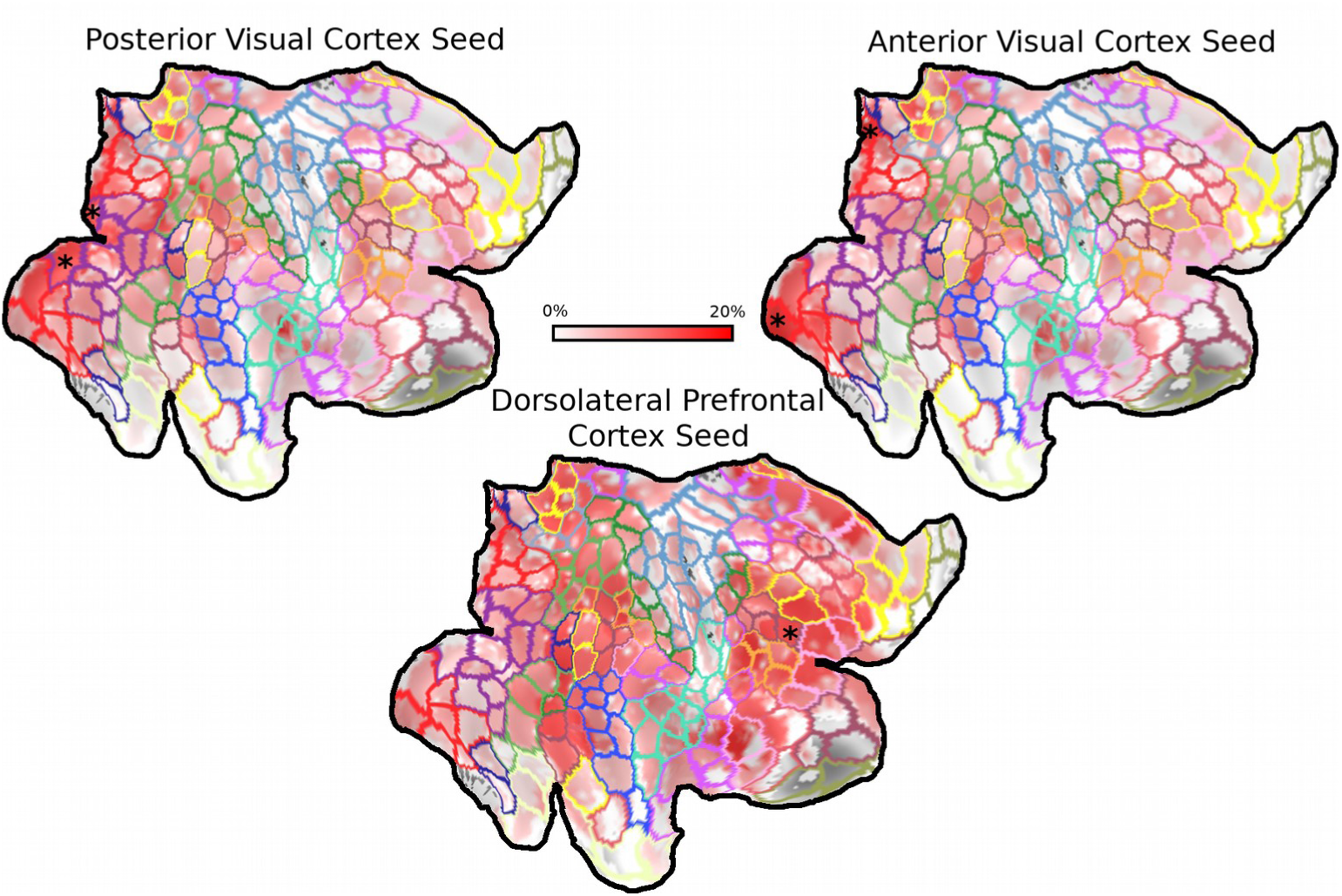
Linearity of input as a general rule of functional connectivity between cortical regions. Mean linearity estimates are shown from 3 seed regions (shown as * in each map), posterior visual cortex, anterior visual cortex, and dorsolateral prefrontal cortex. The color scale from darker to brighter color represents lower to higher linearity for the seed regions’ connective field in the other brain regions. This whole brain illustration demonstrates that while linearity is a general phenomenon for region-to-region connectivity, it is maintained within specific networks, suggesting function in maintaining integrity of information transfer across regions within a network.

To visualize and quantify the network basis of topographic organization of region-to-region connectivity, we constructed a graph representation of our rank linearity maps (Fig. 5A). We then quantified within and between network linearity for each of the 10 MSC subjects across 3 different networks (VisCent, VentralAttenB, DorsalAttenB). For each of the 6 possible comparisons, the within network rank linearity was significantly higher than the between network rank linearity (t’s(9) = 9.52, 3.80, 9.26, 3.19, 9.75, 6.75, p’s = 5.38 x 10^-6^, 4,00 x 10^-3^, 6.76 x 10^-6^, 1.10 x 10^-2^, 4.40 x 10^-6^, 8.33 x 10^-5^) (Fig. 5B).

**Figure 5:**
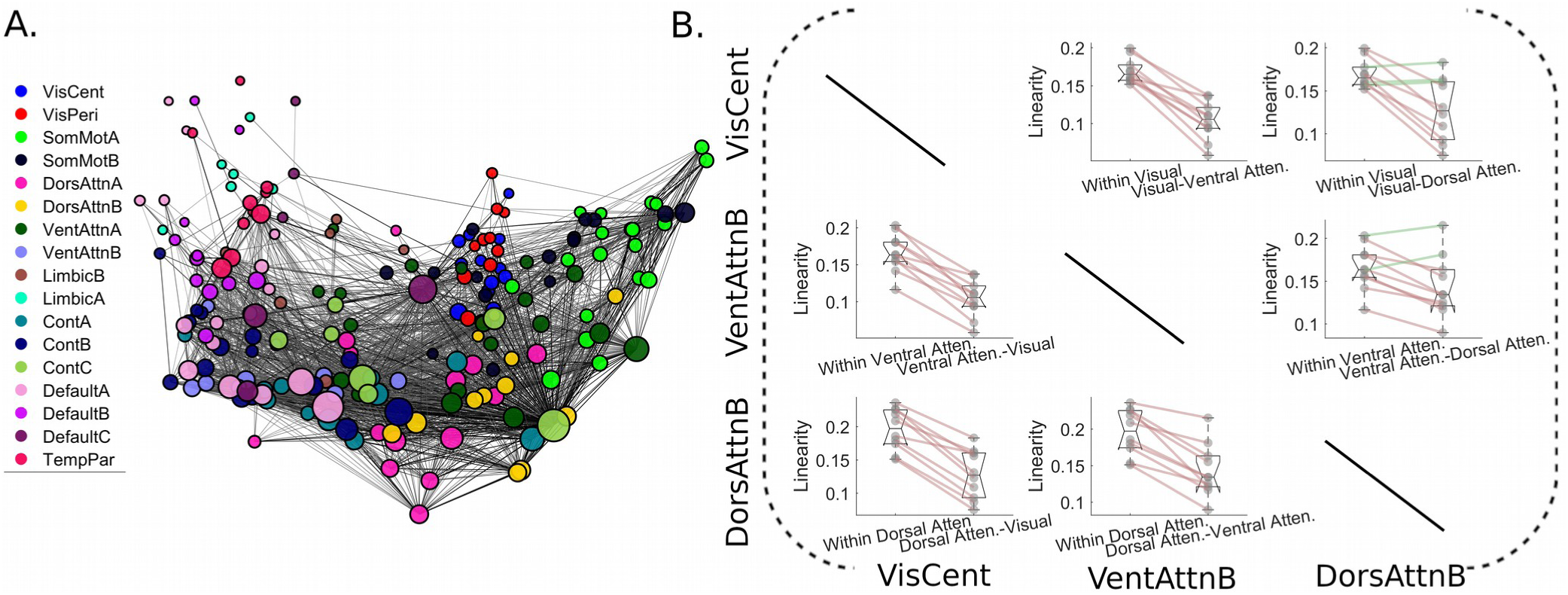
Network Analysis. **(A)** Graph description of our linearity network using the 17 network labels from the Schaefer atlas. Node position was determined by performing classical multidimensional scaling on the full linearity matrix, and then an edge was drawn if it surpassed a linearity threshold of 20%. Node size was scaled linearity in relation to the node’s degree in the thresholded network. **(B)** Quantification of the network like organization of linearity with three example networks, Visual (central), Ventral Attention B, and Dorsal Attention B. The within network linearity is universally higher than the between network linearity in these three networks, as can be seen in **(A)** by the relative closeness of nodes within a network. Note: Schaefer atlas used for network labels, see (Schaefer et al., 2018) for more details.

To examine if there are any higher order (supranetwork) organizational features of topographic connectivity, nonmetric multidimensional scaling was applied to produce two dimensions that maximally reproduce the nonlinearity matrix (as a distance matrix) the 200 parcels in the right hemisphere (Fig. S4A, Fig. 6). The resulting primary mode was similar to large scale whole brain decompositions observed in previous studies of normal functional connectivity and genetic expression data (Margulies et al., 2016; Murray et al., 2018). This mode described a segregation lower order unimodal regions and higher order multimodal regions, with the negative end exemplified by visual and somatomotor networks and the positive end exemplified by default mode network. The second mode seemed to be a fractionation of the primary mode, potentially representing communication of information across these lower order/higher order boundaries.

**Figure 6:**
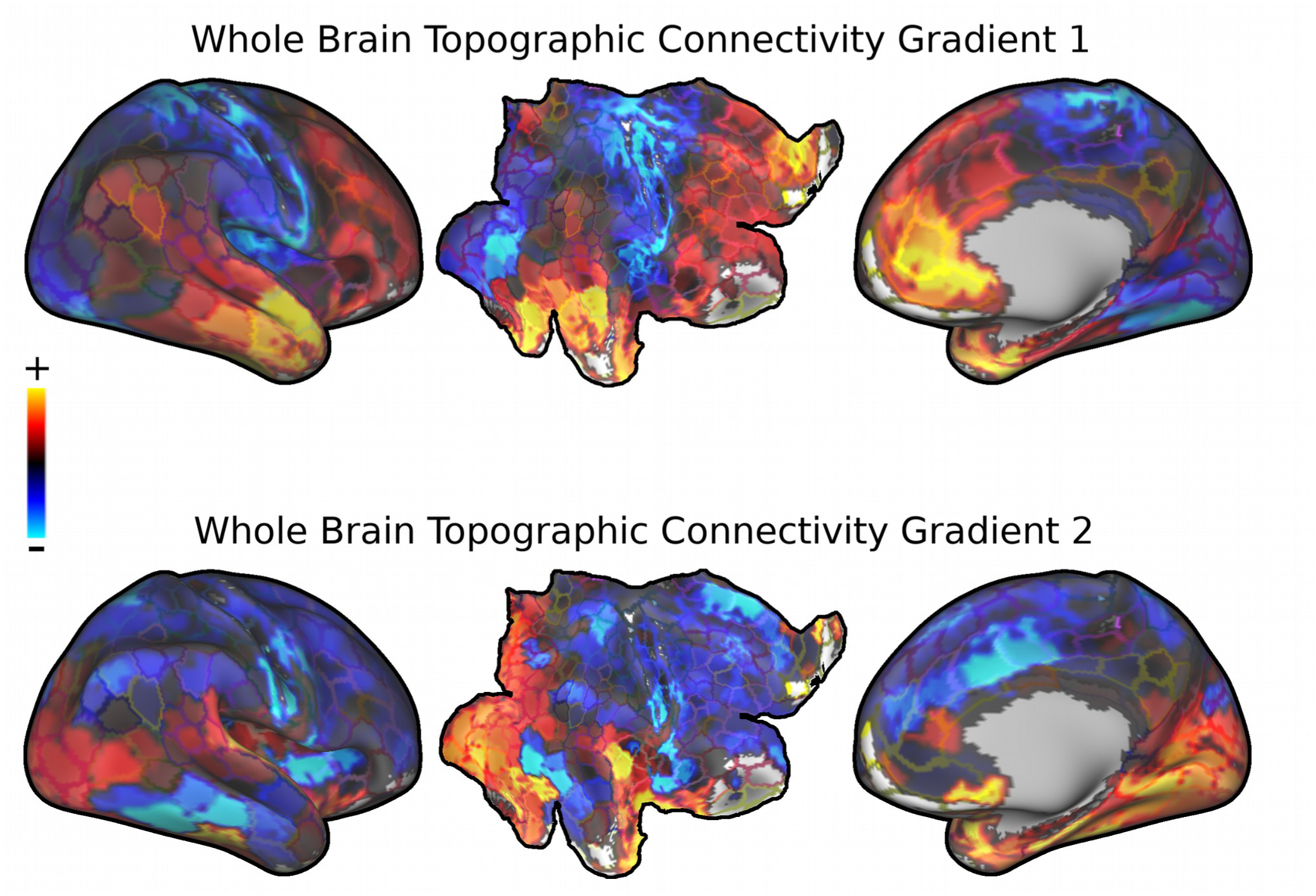
Gradient decomposition of whole brain linearity. Whole brain region-to-region linearity matrix was decomposed using nonmetric multidimensional scaling in order to examine organizational patterns of linearity across the brain. The gradients are color scaled from red to blue, with red-yellow colors representing one side of the gradient, and blue-green colors representing the other side of the gradient. **(TOP)** First whole brain linearity gradient, demonstrates a segregation of unimodal sensory areas and multimodal association areas. **(BOTTOM)** Second whole brain linearity gradient, the fractionated nature suggests potential integration of information across unimodal sensory areas and multimodal association areas.

### 3.4 Connectivity Organization of the Lateral Prefrontal Cortex

After illustrating the ubiquity of linear organization of functional connectivity across visual cortex and large networks, we specifically examined how gradients of linearity express in a highly multimodal structure such as the lateral prefrontal cortex. We found two gradients (Fig. S10B). Gradient 1 demonstrates a rostral-caudal axis that segregates posterior middle frontal gyrus/inferior precentral sulcus and anterior inferior frontal gyrus/frontal pole (Fig. 7A Top Left). Gradient 2 demonstrates a diagonal axis, segregating posterior superior frontal gyrus and anterior inferior frontal gyrus (Fig. 7A Top Right). These two linearity gradients closely resemble the two prominent frameworks described in the literature to simplify the overall functional organization of the frontal cortex. One is often referred to as the domain specialization framework, suggesting that the dorsal and ventral lateral prefrontal cortices are preferentially organized to process spatial and nonspatial (e.g. objects) information, respectively, due to their differential connectivity with the dorsal and ventral visual streams (Goldman-Rakic, 1987). The second is an increase in abstraction along the rostral-caudal axis, with the more rostral regions linked to more abstract facets of behavioral control (e.g. Sensory perception → Context → Rule) (Koechlin et al., 2003). In addition, we performed nonmentric MDS on the standard voxelwise resting-state functional connectivity of the lateral prefrontal cortex, and found two gradient decompositions that mainly segregate middle frontal gyrus and inferior frontal gyrus from the rest of the frontal cortex (Fig. S11).

**Figure 7:**
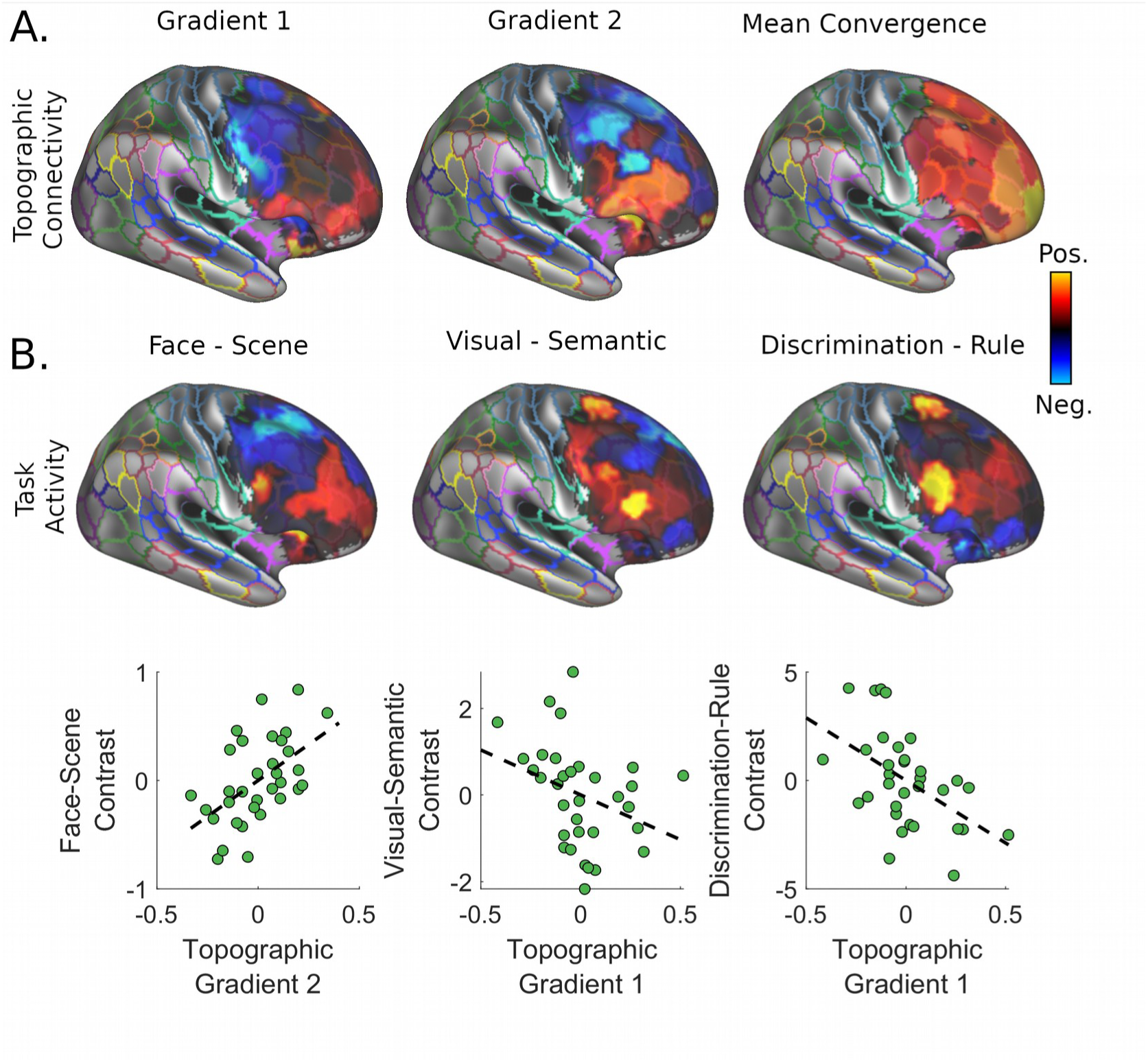
Patterns of functional connectivity within lateral prefrontal cortex. **(A)** The decomposition of the bivariate linearity of the lateral prefrontal cortex revealed two gradients. Gradient 1 demonstrates an organization from posterior middle frontal gyrus/inferior precentral sulcus to anterior inferior frontal gyrus/frontopolar cortex, while Gradient 2 demonstrates an organization from posterior superior frontal cortex to anterior inferior frontal gyrus. The average convergence factor demonstrates a peak along the frontal pole. **(B)** Three different contrasts were examined to correlate to frontal organizational features: face-scene to examine visual specialization, visual-semantic discrimination to examine sensory specialization, and discrimination-rule to examine levels of abstraction. **(C)** First-order correlation demonstrating the most highly associated organizational feature with each contrast. See Table 1 for full regression analysis results.

**Table 1:** Organizational Features of Lateral Frontal Cortex Relates to Task Activation

| Face-Scene Contrast (Incidental Memory Task): $R^2 = 0.37$ | | | | |
| --- | --- | --- | --- | --- |
|  | Beta | Standard Error | T-value | P-value |
| Intercept | 0.00 | 0.14 | 0.00 | 1.0000 |
| Gradient 1 | -0.06 | 0.15 | -0.41 | 0.6822 |
| Gradient 2 | 0.55 | 0.15 | 3.71 | 0.0009 |
| Convergence | 0.31 | 0.15 | 2.08 | 0.0469 |
| Visual-Semantic Contrast (Mixed Task): $R^2 = 0.24$ | | | | |
|  | Beta | Standard Error | T-value | P-value |
| Intercept | 0.00 | 0.16 | 0.00 | 1.0000 |
| Gradient 1 | -0.39 | 0.17 | -2.34 | 0.0265 |
| Gradient 2 | 0.26 | 0.16 | 1.60 | 0.1207 |
| Convergence | 0.26 | 0.17 | 1.56 | 0.1293 |
| Discrimination – Rule Contrast (Mixed Task): $R^2 = 0.41$ | | | | |
|  | Beta | Standard Error | T-value | P-value |
| Intercept | 0.00 | 0.14 | 0.00 | 1.0000 |
| Gradient 1 | -0.56 | 0.15 | -3.83 | 0.0006 |
| Gradient 2 | -0.26 | 0.14 | -1.85 | 0.0751 |
| Convergence | 0.28 | 0.15 | 1.89 | 0.0681 |

We examined whether the average task activation patterns for sensory specialization and level of abstraction from the MSC data sample (Fig. 7B) can be predicted by these organizational features. We found that a linear model with each topographic gradient and convergence predicts each of the activation patterns from each of the contrasts (Face-Scene: R^2^ = 0.37, p = .003 .01; Visual-Semantic: R^2^ = 0.24, p = .046; Discrimination-Rule: R^2^ = 0.41, p = .002). The strongest predictor of Face-Scene contrast was the Gradient 2 value (β = 0.55, p = 8.74 x 10-^4^), while the strongest predictor for the Discrimination-Rule contrast was the Gradient 1 value (β = −0.56, p = 6.34 x 10^-4^) (Fig. 7C, Table 1). This supports the above interpretation of the two gradients of topographic connectivity as relating to domain specialization and abstraction, respectively.

## 4. Discussion

Using a 3-D Gaussian modeling approach with whole-brain resting-state fMRI data, we found that closely connected cortical regions exhibit a relatively strong linear topology of their functional connectivity. This topographic organization of region-to-region functional connectivity was highly reliable in early visual regions, following an order along the anterior-posterior axis for eccentricity representation, as predicted by the literature (Arcaro et al., 2015; Gravel et al., 2014; Haak et al., 2013). In higher order visual regions, such as the FFA and PPA, the estimated connectivity profile seemed to reflect complex filtering of information, such that FFA connectivity to V1 produced the hypothesized patterns of bias towards the part of V1 that has more foveal representations. Further, the PPA shows a more maintained topography with V1 in comparison to PPA. These results are consistent with previous attempts to compare these two higher order regions using a different multivariate method to estimate voxel level patterns of connectivity from one region to another (Baldassano, Iordan, Beck, & Fei-Fei, 2012), where it was found that FFA shows connectivity bias to more foveal resources, while PPA shows a connectivity bias to more eccentric resources. In the whole brain, linearity again emerged as a primary topographic organization for functional connectivity among regions within large-scale networks, including the default mode and frontoparietal networks, while such relationships were much weaker between regions across different networks. Lastly, we found that the topographic and convergence patterns of connectivity in the lateral prefrontal cortex supports the two prevailing frameworks of prefrontal cortex functional organization, domain specialization and abstraction. Our overall findings thus suggest a ubiquitous nature of topography in spatial organization of functional connectivity among closely related functional regions in the human brain.

### 4.1 Eccentricity dependent functional connectivity in early visual cortex as a special case of linear communication

We found that the early visual cortex demonstrates orderly region-to-region functional connectivity that maintains the organization of eccentricity representation across regions. This result replicates previous work that used resting state fMRI data and phase-locking fMRI task data in studies of retinotopic organization of the visual system (Arcaro et al., 2015; Gravel et al., 2014; Haak et al., 2013). The orderly functional connectivity likely contributes to the maintenance of retinotopic organization across the posterior cortical regions for multiple levels of visual processing. We also demonstrated that this motif breaks down for connectivity between early and higher order visual cortices (Fig. 4). The pattern of FFA connectivity demonstrates predicted qualities given its receptive field properties, that is, its convergence and selection of foveal resources (Kanwisher, McDermott, & Chun, 1997). These findings on region-to-region connectivity patterns provide a potential structure for understanding what information is maintained across regions, which would be useful for future modeling of information transformation and transfer from one region to another region (Basti et al., 2018; Thivierge & Marcus, 2007).

The topographic organization of functional connectivity observed in the early visual cortices seems to be a special case of a larger phenomenon. Our findings suggest that such a spatial layout of functional connectivity is a probable general mode of communication across the brain (Fig. 5, 6), likely at least partially for the purposes of high fidelity information transmission across regions within a functional network (Thivierge & Marcus, 2007). In the visual cortex, coincidental activity that drives plasticity provides a potential mechanism for the generation and maintenance a spatially linear organization of communication between nearby regions (Catalano, Chang, & Shatz, 1997; Elliott & Shadbolt, 1996; Schoen, Leutenecker, Kreutzberg, & Singer, 1990). However, activity dependent synaptic pruning is not the only mechanism that drives organization in early visual cortex. Recent studies have suggested a form of proto-organization driven by genetic expression and their products (e.g. morphogens) in animal models of development and in humans (Arcaro & Livingstone, 2017; Cheng, Nakamoto, Bergemann, & Flanagan, 1995; Crowley & Katz, 2000). This mechanism of organization developing from molecular gradients is more widely observed, pertaining to many brain systems. For example, molecular signaling has been shown to be responsible for organization of thalamocortical connections in and ordered spatial gradient, with Emx2 and Pax2 regulating the anterior-posterior organization of cortical fields and their thalamic afferents (Bishop, Goudreau, & O’Leary, 2000). Using 3-D connective field modeling, we found a strong linear organization in lateral prefrontal areas despite a lack of clear or consistent retinotopy in these areas (Hagler & Sereno, 2006; Kastner et al., 2007). Strong linearity in the absence of retinotopy suggests a potential link with proto-organization. Indeed, our gradient decomposition of linearity resembles gradients of genetic expression (Burt et al., 2018; Murray et al., 2018) providing potential convergent evidence for this interpretation.

### 4.2 Topographic connectivity and computation within and across large-scale brain networks

Spatially linear organization of input-output relationships is thought to maintain information across levels of processing, allowing for precise action on continuous sensory information (Thivierge & Marcus, 2007). Topographic organization and projections from topographic space has been postulated to produce ordered abstract space as shown in connectionist models (Tinsley, 2009). With a continuous space of abstract relations, complex information transformation in abstract space can be achieved by the same general mechanism as filtering of visual information. For example, this is the proposed mechanism underlying saliency map models, where information from many independent channels are combined topographically to represent the overall visual salience of an image (Itti & Koch, 2000; Roggeman, Fias, & Verguts, 2010). These saliency map models have been successful in predicting eye movements to complex visual stimuli, attentional load responses in parietal cortex, and even working memory capacity (Foulsham & Underwood, 2008; Knops, Piazza, Sengupta, Eger, & Melcher, 2014).

With the simple assumption information is spatially segregated, a tractable form of general computation in the brain is through spatial biases in connectivity, which would subsume the above description of topographic connectivity. Spatial biases can, however, have vastly different scopes. With distinct information converging onto a neuron giving rise to many potential combinations of that information, it has been shown that the exact combination is at least partially determined by spatial biases in convergence along the dendritic compartment (London & Häusser, 2005; Taylor, He, Levick, & Vaney, 2000). Higher order spatial biases, from cortical column to cortical column, may be responsible for higher order computation. Previous work has examined information transformation more directly, estimating linear transformations that map the representational dissimilarity matrices from one region to another (Basti et al., 2018). Our 3-D connective field model provides an appropriate description of the proposed mechanisms for these transformations, allowing for critical tests of this theory in observational fMRI data.

### 4.3 Organization of the Lateral Prefrontal Cortex Using Spatial Connectivity Metrics

Gradient decomposition of linearity of the prefrontal functional connectivity demonstrated evidence for two gradients of functional organization. Gradient 2, particularly, seemed to predict task based activational differences between face and scene perception. Previous studies have consistently found differential projections from dorsal and ventral visual stream nodes to dorsal and ventral prefrontal cortex, respectively (Cavada & Goldman-Rakic, 1989; Goldman-Rakic, 1987; Kawamura & Naito, 1984). Indeed, scene processing, while involving specialized regions of the ventral visual pathway, does seem to also involve the dorsal stream (Aminoff & Tarr, 2015). This segregation of dorsal and ventral pathway input into the prefrontal cortex has been supported both structurally and functionally in humans (Takahashi, Ohki, & Kim, 2013). This dorsal-ventral segregation of the prefrontal cortex has been shown to be differentially responsible for spatial and object working memory (Constantinidis & Qi, 2018). Indeed, inactivation of dorsal prefrontal cortex impairs spatial working memory, but not object working memory (Chafee & Goldman-Rakic, 2000; Clark, Noudoost, & Moore, 2014; Suzuki & Gottlieb, 2013). Thus, the differential input from dorsal and ventral streams has functional consequences, leading to the segregation of downstream dependence of higher order functions depending on the type of information being operated on.

While a prominent perspective of prefrontal organization, the domain specialization hypothesis of lateral prefrontal functional organization is not the only studied organizational principle.

Accumulating evidence shows another prominent gradient of prefrontal functional organization along the anterior-posterior axis related to the level of abstraction, from the posterior representation of sensorimotor response selection to the more abstract representation of task goals/rules in the anterior portions of the prefrontal cortex (Azuar et al., 2014; Koechlin et al., 2003; Kouneiher, Charron, & Koechlin, 2009). By varying the rules and contexts of various tasks, it was found that areas related to the stimulus response mapping were located more caudally, while areas related to contexts and rules were more rostral (Azuar et al., 2014; Badre & D’Esposito, 2007; Koechlin et al., 2003). While we found that convergence follows a posterior to anterior gradient, there was no evidence that it related to the rule contrast in the mixed task. Patterns of convergence in the lateral prefrontal cortex were, however, associated with different sensory information, likely relating to differential processing demands required. Topographic Gradient 2, on the other hand, did strongly related to the rule contrast in the mixed task, suggesting that topographic connectivity within the lateral prefrontal cortex delineates multiple pathways of organization.

In sum, on the basis of these new metrics of the spatial profile of functional connectivity, our data support both frameworks of functional organization for the lateral prefrontal cortex. However, based on these results it may be the case that these two functionally defined organizations are a result of the same underlying neural organization, driven by the differential topographic input within the lateral frontal cortex. This is consistent with the previous literature examining functional connectivity which has found evidence for both dorsal-ventral and rostral-caudal organizations in the lateral prefrontal cortex (Blumenfeld, Nomura, Gratton, & D’Esposito, 2013; Schumacher, Schumacher, Schelter, & Kaller, 2019).

### 4.4 Limitations and future directions

Our estimations of linearity of connectivity between across regions were likely subjected to parcel selection, which is a limitation for this type of analysis. Given the multitude of parcellations and the multitude of methods used to obtain parcellations, the results could look slightly different if other justified parcellations were used. It is likely that our estimations of linearity would be even stronger with “true” parcels, given that more specification in early visual cortex actually boosted the estimated linearity (from ∼40% to ∼60%). In addition, the 3-D Gaussian model itself also has limitations and makes certain assumptions. For one, it assumes a single focal point of connectivity from one region to another, which is probably overly simplified. However, the final R^2^ of the optimization procedure provides a marker for the fit of the model to the data; if these values are too low in the analysis, it would be inappropriate to make any conclusions about the fitted values. As the R^2^ of the model tended to increase with more scan time, a large proportion of the misfit is likely due to noise. With the 30 minute sessions of the MSC data, the modal R^2^ tended around 0.30-0.60, indicating a reasonable fit. Gaussian mixture models can be developed in the future to account some conditions of misfit, such as regions with multiple maxima.

It is important to note that the 3-D connective field method has a high dependence on high signal-to-noise ratio in single voxels, as opposed to the more conventional analyses that either use signal averaging within ROIs or smoothing to increase the signal-to-noise ratio. Because of this, higher resolution datasets may suffer from less stability due to the increase of proportion of thermal noise in each voxel (Edelstein, Glover, Hardy, & Redington, 1986), though the impact on signal-to-noise ratio can potentially be counterbalanced by a reduction in physiological noise (Bodurka, Ye, Petridou, Murphy, & Bandettini, 2007). Further studies are needed to test our approach on higher resolution data. Optimally, the goal is to achieve a reliable connective field mapping at the size of functional columns of the regions of interest, which may be feasible in the frontal cortex, where columns may be as large as 800-900 micron (Hirata & Sawaguchi, 2008; Masse, Hodnefield, & Freedman, 2017).

### 4.5 Conclusions

By applying 3-D connective field modeling on resting-state fMRI data, we demonstrated that topographic organization of functional connectivity is likely common mode of communication across the cerebral cortex. This pattern of connectivity organization is most evident within the same functional network module, with the whole brain illustrating a primary gradient of linearity of functional connectivity comparable to two different data sources: resting-state functional connectivity and genetic expression. By the same principles, the topographic connectivity of the lateral prefrontal cortex seemed to be organized around two axes: posterior superior frontal gyrus → anterior inferior frontal gyrus and posterior middle frontal gyrus → anterior inferior frontal gyrus / frontopolar cortex. Both these axes segregated spatial patterns of activation relating to domain specialization (Spatial→Object) and abstraction (S-R→Rule). These findings show that deriving finer scale (voxel level) spatial organization of region to region connectivity in volumetric space seems to be useful in resting-state fMRI data. Vector field representations of region to region connectivity has the potential to be an informative visualization for information transfer and transformation from one region to another, with linearity and convergence factor as the quantitative metrics, especially if the voxel size can be reduced to the size of a cortical column in the regions of interest.

