## Supplemental Materials for "Topographic mapping as a basic principle of functional organization for visual and prefrontal functional connectivity"

Supplemental Methods

*Probabilistic Model of Rank Linearity (topography)*

In order to determine whether or not the topographic arrangement of a region's functional connectivity is above and beyond the base topography expected due to chance (given BOLD data characteristics), we formulated a way to examine what it means for a region to be topographically organized. Given the probability of a voxel (in a volume of randomly distributed data) having a data rank order equal to it's spatial rank [p(r)] is the inverse of the volume (V) of the seed:

$$p(r)=\frac{1}{V}$$

Then, the expected rank linearity of a volume is the average probability of this occurrence across the volume:

$$E(Lin)=\frac{\sum^{v} p(r)}{V}=\frac{\sum^{v}\frac{1}{V}}{V}=\frac{1}{V}$$

then, understanding the effect of smoothing on this expected value is understanding the extent to which smoothing reduces the number of independent observations in the spatial field, as defined by the number of resels in the volume:

$$E(Lin_{smoothness})=\frac{1}{V/FWHM^{D}}$$

where FWHM is the estimated smoothness of the volume, and D is the dimensionality of the data (3 in the case of fMRI data). Statistical inferences can be made by evaluating the 95% confidence intervals of this probability on a binomal distribution given the number of voxels in the volume.

To examine whether the process described actually reflects the behavior of our rank linearity estimations, we simulated volumes of randomly distributed data while manipulating both the number of voxels in the volume and the smoothness. First, we showed that the expected value of rank linearity is approximately equal to the probability of a rank, which is the inverse of the voxels in a volume (Fig. S1A). Secondly, we found that the effect of smoothing follows from the approximate number of independent observations (i.e. number of resels) within the volume (with the 95% confidence intervals well approximated by evaluation on the binomial) (Fig. S1B). However, we noted that there seemed to be some divergence of the model from the simulated data at higher smoothing values. We then conducted more simulations to test if the deviance was related to the total number of independent observations in the volume, by manipulating the number of voxels in the volume. If the deviance is due to the total number of independent observations (resels) then this data should diverge from the model with a lower smoothing FWHM in simulations with fewer voxels. We found that this was roughly the case (Fig. S2B (volume = 125 voxels) vs Fig. S2A (volume = 729 voxels)). We selected a rule of thumb to flag inferences that may be inappropriate, where the number of resels in a ROI is less than or equal to 10. None of the analyses in the main manuscript were flagged.

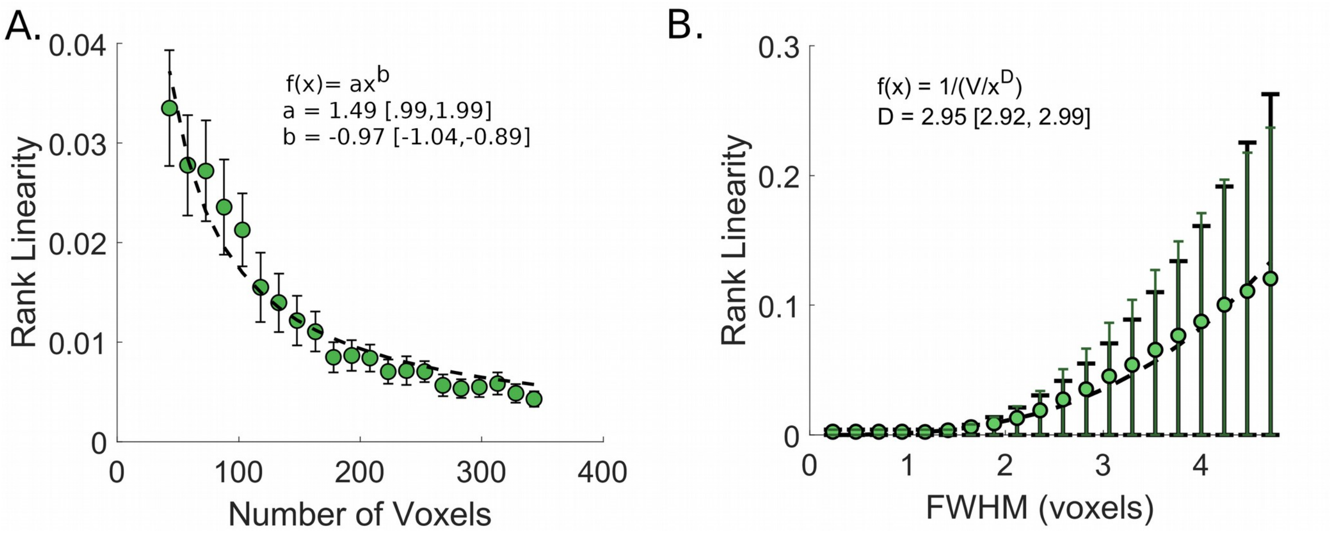


Figure S1: Simulations results of rank linearity and probabilistic model of rank linearity. **(A)** Rank linearity approximates the probability of a given rank value. Model parameters represent best fit values obtained by nonlinear least squares fitting in Matlab's curve fitting toolbox. **(B)** Effect of data smoothness on rank linearity is related to the estimated number of resels in a region. 95% confidence intervals of the simulation and model are displayed as errorbars on each point. Simulation values in green dots and model values in black dashed lines.

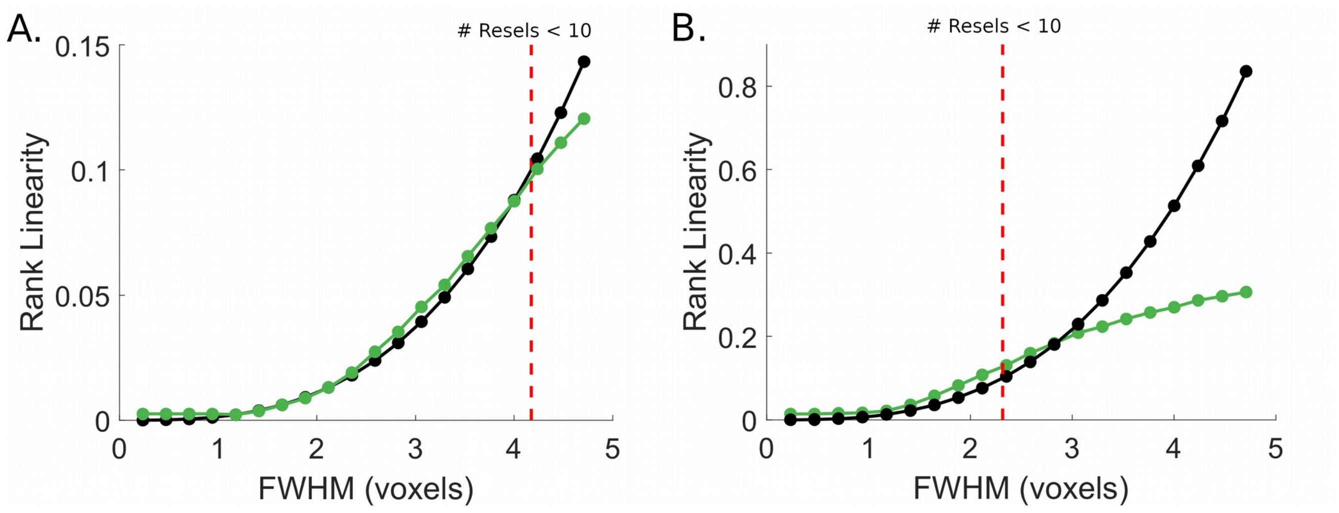


Figure S2: Simulations (green) deviate from model data (black) at low number of resels as shown by two examples: **(A)** a region of 729 voxels, and **(B)** a region of 125 voxels. We reasoned that if this was a result of number of total resels in a region, then this deviance should occur at a lower smoothness value in regions with a lower number of voxels. This is what we observed comparing **A** and **B**, As a rule of thumb, we tag all calculations of rank linearity when the number of resels are below 10. While no observations in the main manuscript were tagged, theoretically one could perform Monte Carlo simulations to determine appropriate error rates in those cases.

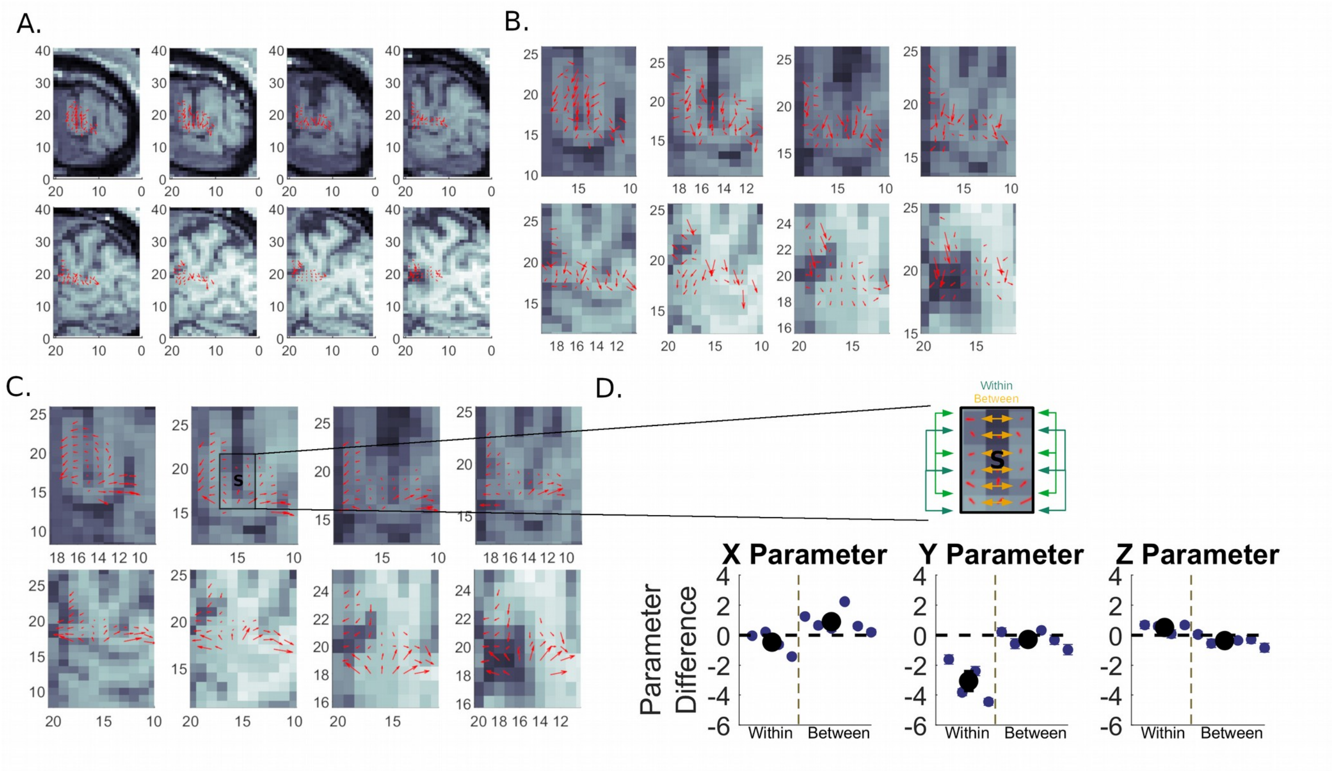


Figure S3: **(A)** Depiction of a single subject's vector field from V1d to V1v plottted on that subject's normalized anatomy. **(B)** Zoomed in version of the same subject's vector field shown in **(A)**. **(C)** An average of 300 subject's vector fields, accentuation the features seen in the vector field in **(B)**. **(D)** Comparison of a voxels across and within sulcus walls. While both X and Z parameters are confounded by space, the Y parameter comparison is not, demonstrating a difference between voxels along a sulcus wall and voxels between sulcus walls.

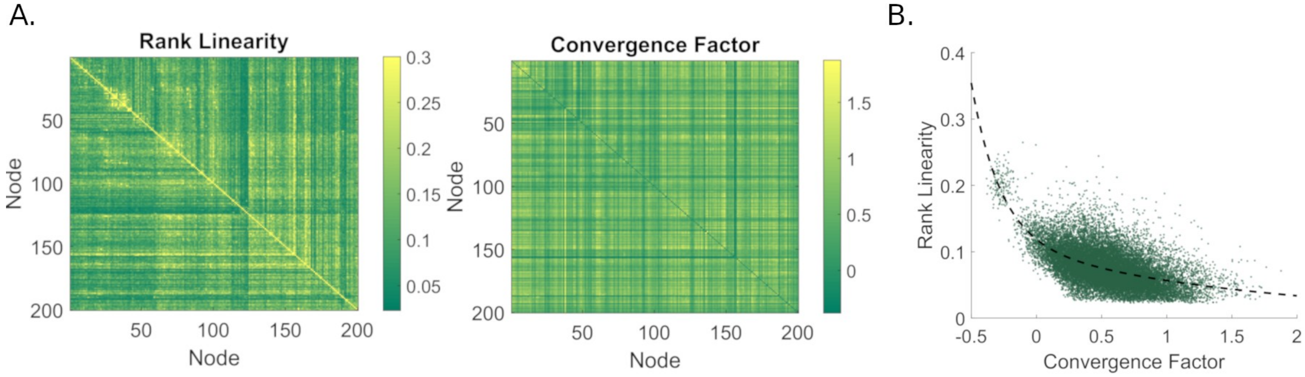


Figure S4: Relationship between linearity and convergence factor. **(A)** The node by node linearity and convergence matrices. An inverse relationship between these metrics can be seen in the pattern of node by node relationships. **(B)** A quantitative comparison of all unique pairs of regions. There is a negative non-linear relationship between linearity and convergence, though this relationship only accounts for a minority of variance (37.06%) across the two metrics.

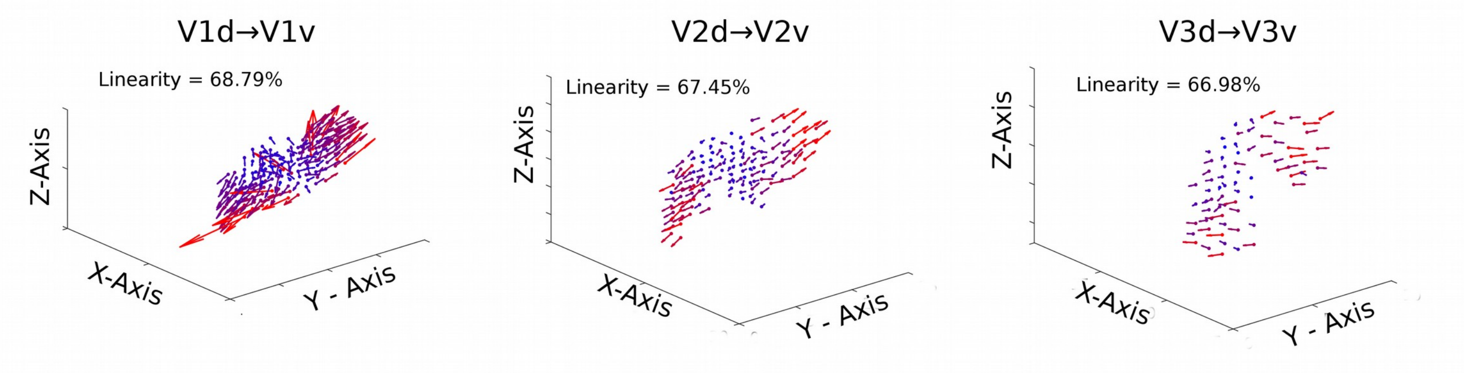


Figure S5: The linear organization pattern of region-to-region functional connectivity is similar across early visual areas. More constrained ROI masks were selected to demonstrate that the connectivity patterns from dorsal to ventral portions of V1, V2, and V3 are constrained by the anterior-posterior axis, likely along the topographic eccentricity representation. The vector fields are displayed similarly as described in Figure 2B, with the red-blue color scale representing the euclidean norm of each seed voxel's vector.

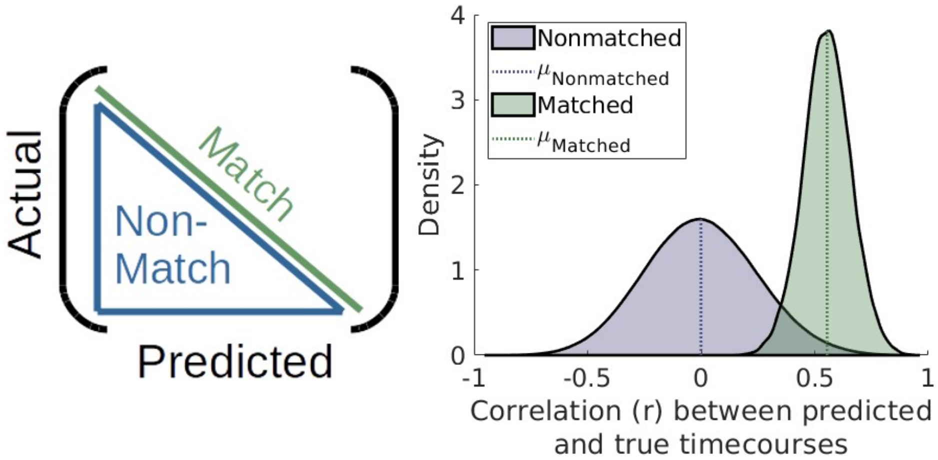


Figure S6: Evaluation of the prediction of each seed voxel's timecourse from the 3-D connective field model. Correlation of the actual timecourses of voxels in the seed region (dorsal visual cortex) with their corresponding predicted timecourses was compared to their correlation and all other predicted timecourses. The distribution matched pairs of correlations is much tighter and shifted positively in comparison to that of the mismatched correlations.

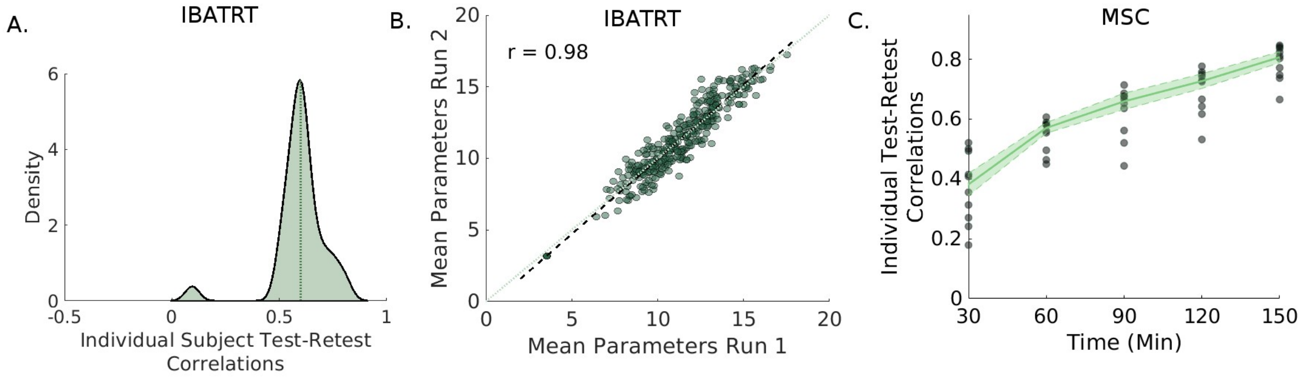


Figure S7: Test-retest reliability examined using the IBATRT and MSC datasets. **(A)** Correlation of each subject's parameters for the V1d seed voxels in the first and second session. **(B)** Correlation between the mean parameters for each of the sessions, the black dashed line is the least squares fit, while the light green dashed line is the identity line. **(C)** The correlation between each MSC subject's parameters for V1d seed voxels while incorporating different amount of data into each test-retest set, maxing out at 150 minutes of data within each set. With 150 minutes of data, the median individual subject test-retest correlation is 0.81.

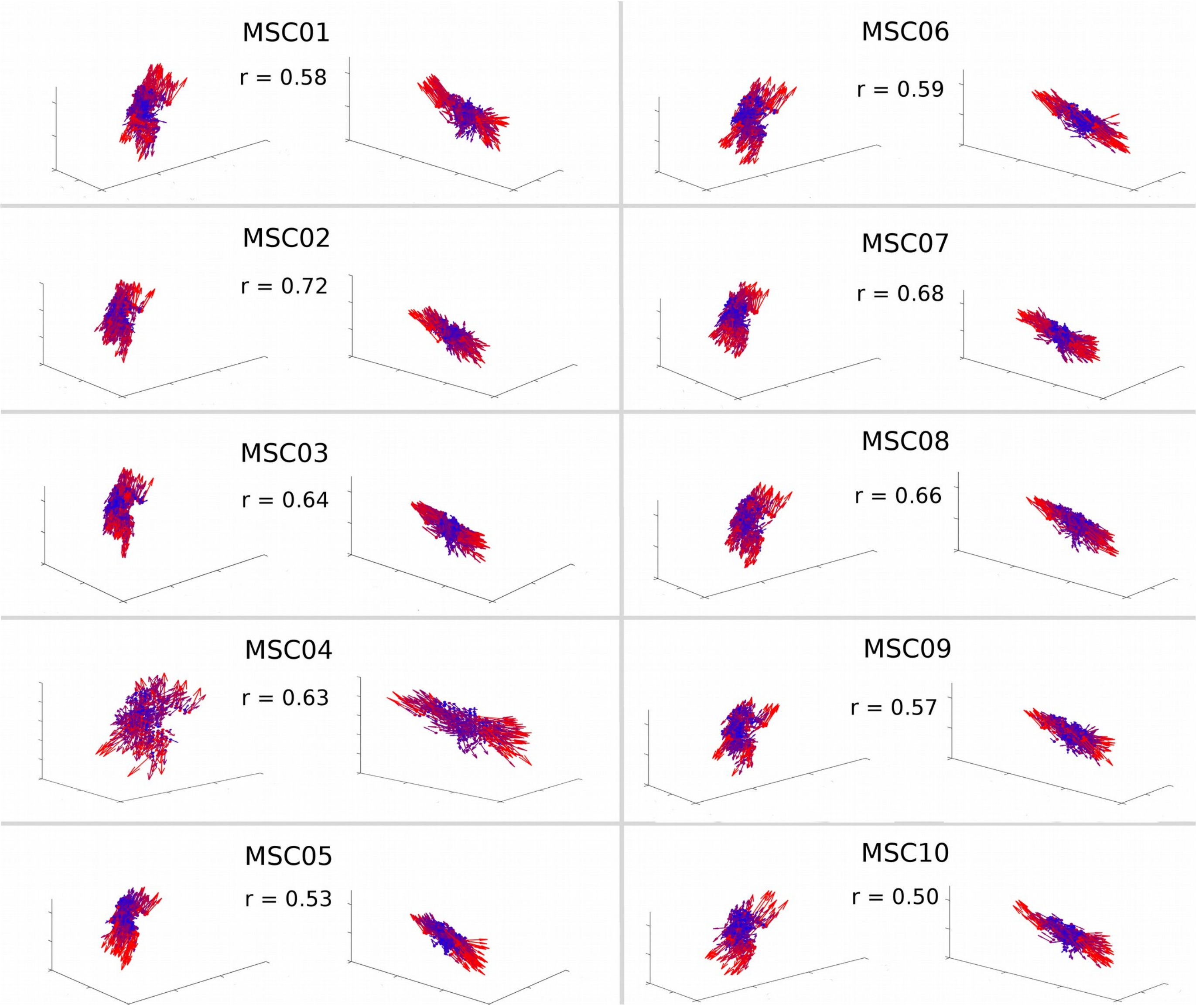


Figure S8: Vector fields from individual subjects in the MSC dataset. The anterior posterior topographic organization of the vector fields is detectable in every subject, as denoted by the r-value in each subject's vector field plots showing the correlation between the y-location parameters and the y-space.

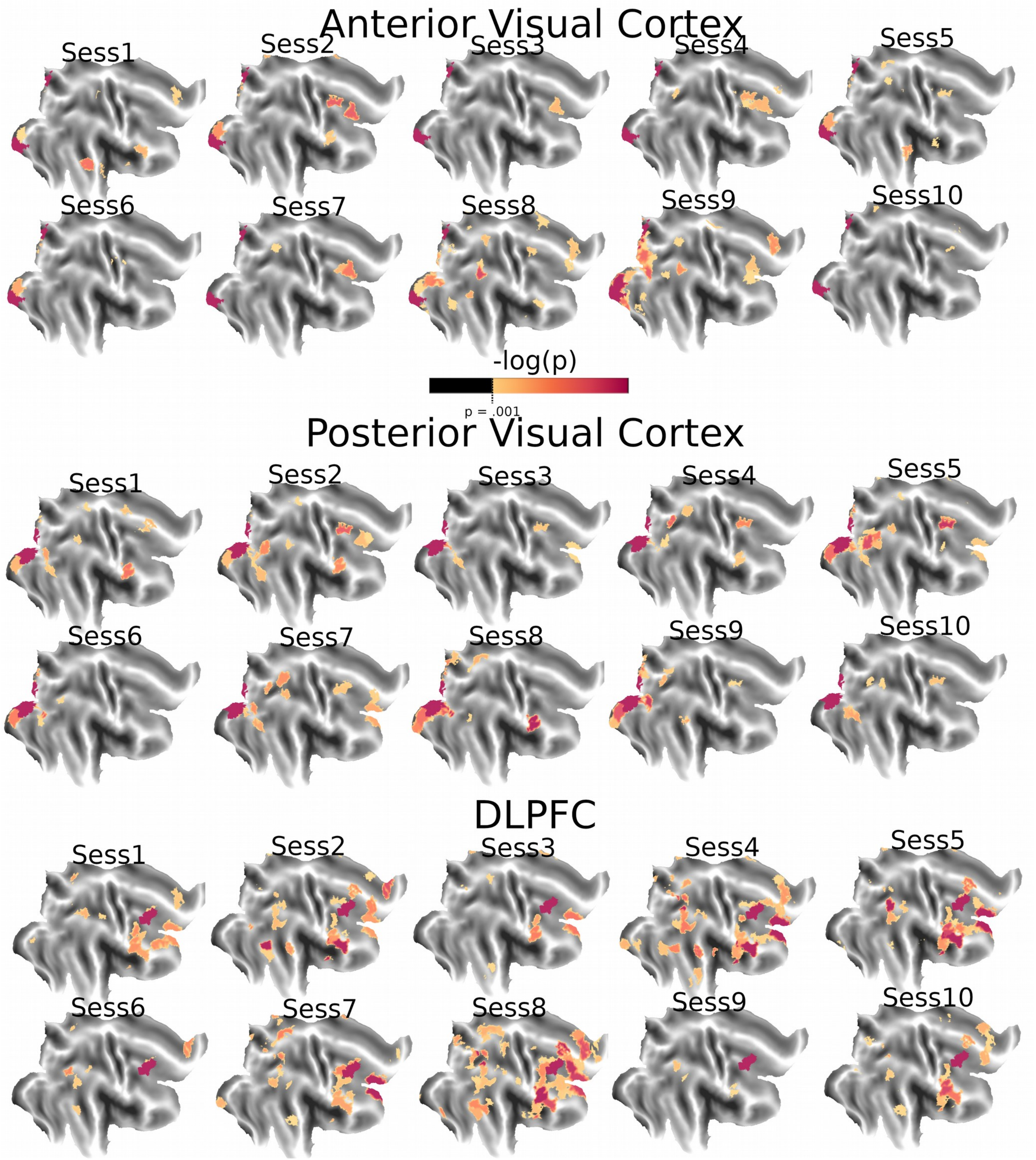


Figure S9: MSC subject 01's seeded topographic maps thresholded at p < .001 for each session. Color maps are scaled on the basis of -log(p) values. **(Top)** Topographic maps of functional connectivity seeded from anterior visual cortex, **(Middle)** posterior visual cortex, and **(Bottom)** DLPFC all demonstrate consistent patterns of linearity within each of their networks across each session. Maps are overlaid on a flatmap aligned with the HCP fslr32k atlas.

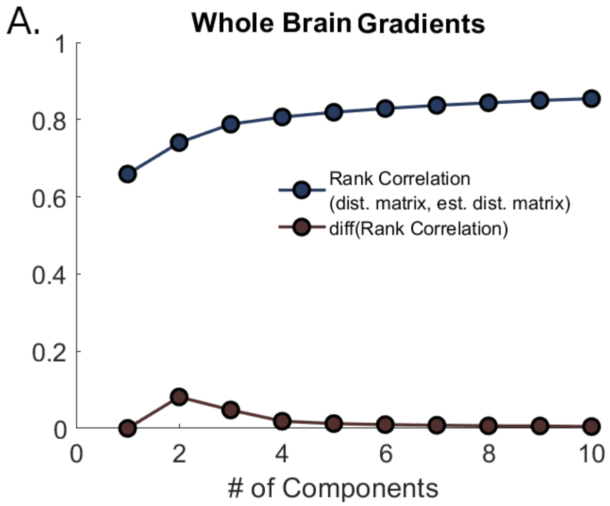


Figure S10: Determination of the number of gradients in the linearity decomposition for the whole brain **(A)** and the lateral frontal cortex **(B)**. We examined the rank correlation between the distance matrix and the MDS estimated distance matrix, and visually identified an elbow by examining where the largest spike lies in the derivative of the correlations with respect to component number. In both cases, this ended up being 2 components.

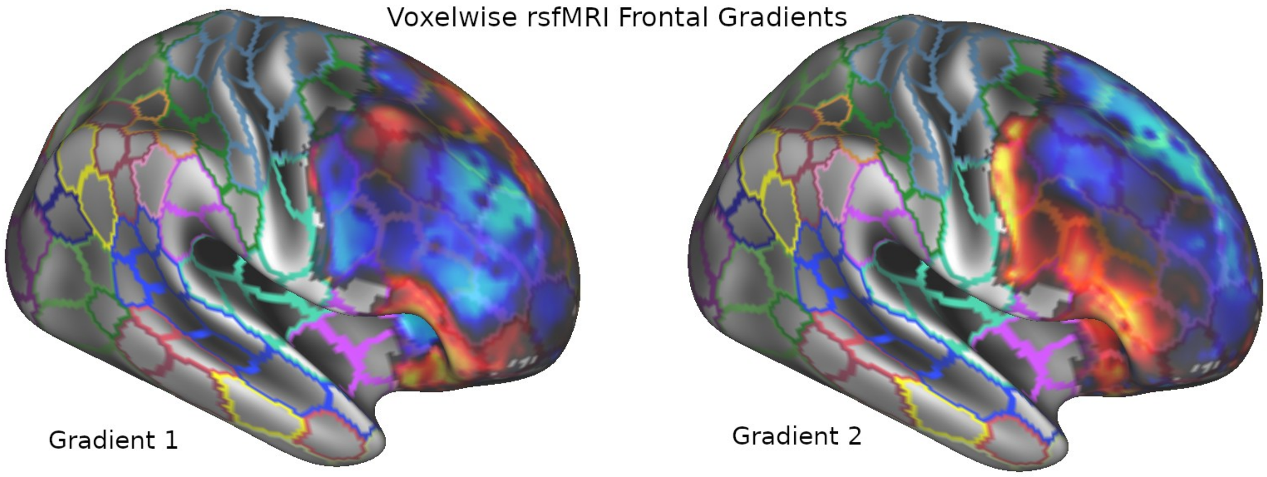


Figure S11: Lateral Frontal gradients estimated from voxel-wise resting-state fMRI data. Gradient 1 segregates Middle Frontal Gyrus from the rest of lateral frontal cortex, while Gradient 2 segregates Inferior Frontal Gyrus from the rest of the lateral frontal cortex. These derived gradients are qualitatively different than the gradients derived from the topographic gradient estimates, demonstrating that these metrics provide unique information beyond the typical resting-state analyses.
